# Free-living psychrophilic bacteria of the genus *Psychrobacter* are descendants of pathobionts

**DOI:** 10.1101/2020.10.23.352302

**Authors:** Daphne K. Welter, Albane Ruaud, Zachariah M. Henseler, Hannah N. De Jong, Peter van Coeverden de Groot, Johan Michaux, Linda Gormezano, Jillian L. Waters, Nicholas D. Youngblut, Ruth E. Ley

## Abstract

Host-adapted microbiota are generally thought to have evolved from free-living ancestors. This process is in principle reversible, but examples are few. The genus *Psychrobacter* (family *Moraxellaceae*, phylum *Gamma-Proteobacteria*) includes species inhabiting diverse and mostly polar environments, such as sea ice and marine animals. To probe *Psychrobacter’s* evolutionary history, we analyzed 85 *Psychrobacter* strains by comparative genomics and phenotyping under 24 different growth conditions. Genome-based phylogeny shows *Psychrobacter* are derived from *Moraxella*, which are warm-adapted pathobionts. *Psychrobacter* strains form two ecotypes based on growth temperature: flexible (FE, growth at 4 - 37°C), and restricted (RE, 4 - 25°C). FE strains, which can be either phylogenetically basal or derived, have smaller genomes and higher transposon copy numbers. RE strains have larger genomes, and show genomic adaptations towards a psychrophilic lifestyle and are phylogenetically derived only. We then assessed *Psychrobacter* abundance in 86 mostly wild polar bear stools and tested persistence of select strains in germfree mice. *Psychrobacter* (both FE and RE) was enriched in stool of polar bears feeding on mammals, but only FE strains persisted in germfree mice. Together these results indicate growth at 37°C is ancestral in *Psychrobacter*, lost in many derived species, and likely necessary to colonize the mammalian gut.

## Introduction

Whether microbiota are associated with vertebrate hosts or not is the largest factor driving differences in the composition of microbiomes sampled globally [1, 2]. Recent analysis of metagenome-assembled genomes from multiple habitats shows that many of these genomes are either animal host-enriched or environment-enriched, but generally not both [3]. Strikingly, such specialization can also be seen at higher taxonomic levels, indicating that whole lineages may have diverged once animal hosts were first successfully colonized. For instance, within the *Bacteroidetes*, the taxa that are mammal-gut associated are derived from phylogenetically basal clades that include free-living and invertebrate-associated taxa [4]. These patterns of distribution imply that specialization to the warm animal host habitat is mostly incompatible with fitness in other environments.

There are known exceptions: microbiota with complex lifestyles adapted to life both on and off the warm animal host. A few taxa from the phylum *Proteobacteria*, many of which are pathogens and pathobionts, inhabit mammalian bodies and have environmental reservoirs [5, 6]. Genomic adaptations that support fitness across several different environment types have been identified, many of which increase infectivity in mammals. For instance, genes that allow bacteria to avoid predation by protozoa, amoebas, and nematodes, also contribute to virulence in mammalian infections in species such as *Vibrio cholera, Burkholderia pseudomallei*, and *Yersinia pestis* [7, 8]. Genes regulating the formation of biofilms have also been implicated in the infectivity of organisms such as *V. cholera* [9]. Type IV pili, organelles that are important for the asymptomatic colonization of plant tissues, are also associated with mammalian tissue invasion [10]. Particular strains of virulent *Escherichia coli* serotype O157H7 show increased or decreased ability to persist in soil depending on mutations in their stress response genes, which impact survival in acid and other selection pressures [11]. In the food-borne pathogen *Listeria monocytogenes*, mutations in genes important for cell invasion also affect cold tolerance, such that strains with increased persistence in processed foods are more likely to have high virulence [12]. For the majority of nonpathogenic animal-associated microbiota, the adaptations to life in and on the host seem to preclude sustaining populations outside the host.

The evolutionary history of pathogens has been studied in depth in a few cases, and indicate an environmental ancestry. The evolutionary trajectory of commensal microbiota is less well characterized, but many likely follow the same patterns. *Mycobacterium tuberculosis* and *Y. pestis* are both pathogens thought to be derived from non-pathogenic, environmental organisms [13, 14]. Many *Mycobacteria* are soil microbes that can sometimes cause disease in mammals, while *M. tuberculosis* itself is only found in humans and has no known environmental reservoir [14, 15]. *Y. pestis* is closely related to pathogenic species *Y. pseudotuberculosis* and *Y. enterocolitica*, though other *Yersinia* spp. are nonpathogenic soil and water microbes [13, 16]. There may be cases of the inverse: where environmental bacteria are derived from animal host-associated ancestors, but well characterized examples are lacking. Experimental evolution studies in *Pseudomonas aeruginosa* and *Serratia marcescens* have established that a trajectory from host to environment is possible, however, and results in attenuated virulence [17, 18]. The more bacterial genomes become available for comparative studies, the better the understanding will be of how certain lineages may have moved between animal hosts and their environments.

Here, we investigated the evolutionary history of the genus *Psychrobacter*, a group of closely related bacteria with a broad environmental distribution. Species of *Psychrobacter* have been recovered through culture-based and sequenced-based methods from a range of animal microbiomes, including marine mammal skin [19], respiratory blow [20], and guts [21–23]; the gastrointestinal tracts of birds [24, 25] and fish [26]; and many nonhost environments such as sea water [27], sea ice [28], marine sediment [29], glacial ice [30], and permafrost soil [31]. Some *Psychrobacter* species are capable of causing disease in mammalian hosts [32, 33]. However, *Psychrobacter* infections are very rare, and the virulence factors involved are relatively uncharacterized. Intriguingly, a previous comparative genomics analysis of 26 *Psychrobacter* spp. and metadata gleaned from public sources revealed differences in cold-adaptation of protein coding sequences between warm-host-associated strains versus derived marine and terrestrial strains [34]. Furthermore, warm-adapted strains were basal in the *rpoB* gene phylogeny, suggesting that *Psychrobacter* evolved from a mesophilic ancestor. These observations make *Psychrobacter* an interesting candidate to assess how ecotype maps onto phylogeny and source of isolation, and to probe into the evolutionary history of a genus with a wide habitat range.

We tripled the collection of *Psychrobacter* genomes, which we use for phylogenomic analysis, and combine these data with extensive phenotyping applied consistently for all strains. We use a large collection of wild polar bear feces collected on ice and land to assess the presence of *Psychrobacter* ecotypes as a function of diet determined by cytochrome b barcode sequencing. Finally, we conducted tests with select strains for colonization of the mammal gut using germ-free mice. Our results confirm a mesophilic ancestry for *Psychrobacter* and a common ancestor with the genus *Moraxella*. Our phenotyping revealed that overall, *Psychrobacter* tolerate a wide range of salinity, but growth at 37 °C divided the accessions into two ecotypes: those that retained the ability to grow at warm temperatures and colonize mammalian hosts (flexible ecotype, FE), and those that exhibit adaptive evolution towards a psychrophilic lifestyle (restricted ecotype, RE). Genomic analysis of the two ecotypes shows genome reduction in the FE strains with high transposon copy numbers, and adaptation to cold in RE strains. We show that *Psychrobacter* that are basal are FE, but that FE are also interspersed with RE, indicating either re-adaptation to the animal host or retention of the basal traits. Although both FE and RE ecotypes were detected in the feces of wild polar bears, only FE strains tested could colonize the germfree mouse gut. Together our results indicate the evolutionary history of the genus *Psychrobacter* indicates a pathobiont losing its ability to associate with animals in its adaptation to nonhost environments.

## Materials and Methods

All software versions, parameters, and the relevant citations for our analyses are detailed in Table S1.

### *Moraxellaceae* family genomics and phenotypic data

The *Moraxallaceae* family includes three well characterized genera: *Moraxella, Acinetobacter* and *Psychrobacter*. To build a family-level phylogeny, we downloaded genomes of 18 species of *Acinetobacter* and 18 species of *Moraxella* from NCBI (June 2020) (Table S2). We also included 15 *Psychrobacter* genomes generated in this study (described below). We determined the phylogenetic relationship between the genomes using whole-genome marker gene analysis software PhyloPhlAn, and determined genome quality and summary characteristics using checkM and Prokka. The *Moraxellaceae* phylogenetic tree was visualized and annotated using the interactive Tree of Life (iTOL) web interface.

We analyzed the *Moraxellaceae* pan-genome using the PanX pipeline. The input genomes used by PanX were initially annotated using Prokka. After their assignment into orthologous clusters using MCL, we re-annotated gene clusters using eggNOG mapper. We explored genome content by calculating a distance matrix using the Jaccard metric through the R package ecodist from a binary gene presence-absence table, followed by dimensional reduction of that distance matrix through principle coordinate decomposition (PCoA) with the cmdscale function from the R package stats. We investigated variables contributing to the separation of the PCoA using the envfit function from the R package vegan; we tested whether genes significantly contributing to separation were “core” (genes present in 90% to 100% of strains), “shell” (genes present in greater than two strains, but in fewer than 90%), or “cloud” (genes present in only one strain), and if general gene function - summarized by Cluster of Orthologous Groups (COG) category - contributed to the separation. We collected growth temperature range data from type strain publications [23, 24, 27, 28, 35–76]. We used the R package phytools [77] to map the temperature ranges onto the phylogeny.

### *Psychrobacter* strains

We obtained 92 isolates of *Psychrobacter* from strain catalogues for phenotypic and genotypic characterization. These represent 38 validly published species of *Psychrobacter* as well as unclassified strains, all isolated from a wide variety of geographical locations and diverse environmental and host samples. All strains were purchased and maintained in compliance with the Nagoya Protocol on Access to Genetic Resources and the Fair and Equitable Sharing of Benefits Arising from their Utilization to the Convention on Biological Diversity. For a full list of accessions and their catalogue, isolation, and cultivation information, see Table S3. Unless mentioned, we grew accessions as recommended by the strain catalogue (medium and temperature) from which they were purchased. We performed electron microscopy as described in the Supplemental Text.

### *Psychrobacter* phenotypic screen

For the 85 *Psychrobacter* accessions that passed genome quality control (see Supplemental Text, Table S4), we tested their ability to grow under 24 different conditions; a growth condition being a combination of a medium (complex or defined), salt concentration (0, 2.5, 5 or 10% NaCl) and incubation temperature (4, 25 or 37 °C). For more information regarding the media choice, see Supplemental Text.

We randomly assigned *Psychrobacter* accessions to blocks of ten strains to be tested simultaneously (several accessions were included in multiple blocks, see Table S2). We first grew strains to saturation, washed, diluted to optical density at 600 nm (OD_600_) = 0.3 in sterile phosphate buffered saline (PBS), and inoculated in 100 μL of medium with a final ratio of 1:1000. We used 96-well plates, with 10 inoculum in 5 replicates and 10 uninoculated media wells per plate, all periphery wells were filled with water to reduce edge effects. Plates were incubated at each temperature to reflect all 24 growth conditions. We measured the OD_600_ (Spark^®^□ plate reader, Tecan, Zürich, Switzerland) every eight hours during the first week of incubation, then every twenty-four hours, and let cultures grow until stationary phase (3 to 12 weeks). We tested the carbon utilization of a subset of *Psychrobacter* accessions as described in the Supplemental Text.

### Growth probabilities

We scored each replicate as either ‘growth positive’, meaning the accession grew, or else, ‘growth negative,’ if the replicate never reached a maximum OD_600_ of 0.15 over the course of the experiment. For each strain and condition (medium, salt and temperature), using base R functions we calculated a growth probability, which corresponds to the median value of growth positivity/negativity of all replicates for that condition.

### Genome sequencing, assembly and annotation

Genomic DNA was extracted from cultures grown in their preferred conditions using the Gentra Puregene Tissue Kit (Qiagen, Valencia, CA, USA). Samples were sequenced using the MiSeq 2×250 bp and HiSeq 2×150 bp paired-end read technology (Illumina, San Diego, CA, USA) as previously described [78], with some samples undergoing additional long-read sequencing using Ligation Sequencing (Oxford Nanopore, Oxford, UK). For details regarding quality control and assembly, see Supplemental Text, and for a summary of genome quality, see Table S4. Note that of the initial 92 accessions, 85 had genomes that passed all QC measures (Table S4). All further analyses use these 85 accessions.

We annotated genomes with Prokka and eggNOG mapper. A phylogeny of the accessions was generated using PhyloPhlAn, with *Moraxella lincolnii* as an outgroup. Again the phylogeny was visualized and annotated using iTOL. We used PanX to analyze the *Psychrobacter* pan-genome and R to explore gene presence-absence data as described above with the Moraxellaceae family. Pseudogenes were predicted using the DFAST core workflow. We used treeWAS to perform a pan-genome wide association study relating data collected in the phenotypic screen to genomic data.

### Potential cold-adaptive trait analysis

For every gene cluster of the *Psychrobacter* pan-genome obtained, we used DIAMOND to map the consensus sequence against the UniRef90 database, excluding results from the family *Moraxellaceae*. We thus obtained homologous sequences while ensuring that the analysis would not be biased by self-comparison. Gene clusters without UniRef90 homologs were removed from further analysis.

The putative cold adaptive traits of every protein-coding gene from the *Psychrobacter* set and their UniRef90 homologs were evaluated following the methods of Bakermans [34]. Briefly, we used the relative abundance of every amino acid per coding sequence (CDS) to calculate the arginine to lysine ratio, acidity, hydrophobicity and KVYWREP ratios [34]. We additionally calculated the grand average of hydropathicity (GRAVY) [79] and the isoelectric point (using Protein Analysis method class from Biopython). Next, we compared the distributions of these measures from *Psychrobacter* protein sequences to the distributions of the measures from the UniRef sequences. For each protein sequence, if a majority of measures were in the top 25% of the homologs distribution, we defined the protein sequence as “highly cold adaptive”. The top 25% were defined as follows for each trait: highest 25% for glycine relative abundance, acidity, hydrophobicity, and GRAVY; and the lowest 25% for proline relative abundance, arginine relative abundance, arginine-lysine ratio, isoelectric point, and KVYWREP.

### Microbiome diversity of polar bear feces

We collected 86 polar bear fecal samples from several regions in Canada, including samples from 10 captive bears, fed varying diets or fasted, and 76 samples from an unknown number of wild bears, whose diets were determined from stool analysis (below). The feces from the captive bears were forwarded from institutions within Canada and did not require permitting for their passage to Queen’s University. The captive samples comprised: five fecal samples from a single bear sequentially fed varying diets of Arctic char, harp seal and a “zoo diet” at the Polar Bear Habitat in Cochrane during 2010; two samples from each of two bears held at the Metro Toronto Zoo, fed a consistent “zoo diet” during 2010; and a single sample from a bear held at the Churchill Polar Bear Holding Facility in Churchill during 2010, where the bears are given only water until release. We collected all wild bear faeces from M’Clintock Channel and Hudson Strait in Nunavut in accordance with permits prior to shipping to Queen’s University: we collected 24 samples from M’Clintock Channel under *Wildlife Research* permits issued in 2007, 2008, 2009, 2010 and 2011 to Peter Van Coeverden de Groot, and nine samples from Hudson Straight collected in 2011 under *Wildlife Research* permit to Grant Gilchrest (Environment Canada). We collected 43 wild samples from the Wapusk National Park in Manitoba from 2007-2010 under a Canada Parks permit to Robert Rockwell (American Museum of Natural History).

We confirmed that samples collected from wild bears originated from bears by sequencing a cytochrome b gene fragment using metabarcoding approaches based on Ion Torrent (Ion Torrent Systems Inc., Gilford, NH, USA) and 454 pyrosequencing (454 Life Sciences, Branford, CT, USA) next-generation sequencing technologies [80]. We also used this method to analyze what prey animals the polar bears had been feeding on at the time of sample deposition. We visually inspected the samples for confirmation on the dietary analysis. We extracted DNA from the fecal samples using the DNeasy Blood and Tissue kit (Qiagen), and characterized the gut bacterial community by amplification and sequencing of the V4 region of the 16S rRNA gene as described previously [81]. We used QIIME for sequence processing and Silva to assign taxonomy. As described in Supplemental Text, we isolated a strain of *Psychrobacter (P. faecalis PBFP-1*) from polar bear feces that we included in our genomic and phenotypic analyses.

### Gnotobiotic mouse colonizations

Based on their source of derivation and phylogenetic breadth, we selected eight accessions of *Psychrobacter - P. cibarius JG-220, P. ciconiae, P. faecalis PBFP-1, P. immobilis S3, P. lutiphocae, P. okhotskensis MD17, P. namhaensis*, and *P. pacificensis* – to monocolonize germfree mice. We grew each accession in its preferred conditions (Table S3) to saturation, spun at 10 °C at 2500 rpm for 25 minutes, washed with sterile PBS, resuspended in 15% (v/v) glycerol in PBS, and flash frozen in liquid N_2_. Inoculum samples were stored at −80 °C until the mouse experiments were performed by animal caretakers as follows. For *P. ciconiae*, *P. faecalis*, *P. namhaensis*, and *P. pacificensis*, experiments were performed at Taconic Biosciences (Rensselaer, NY, USA), while for *P. cibarius*, *P. immobilis*, and *P. okhotskensis*, experiments were performed at the Max Planck Institute for Developmental Biology. *P. lutiphocae* was included in both Taconic and MPIDB experiments.

Five to six-week old germfree male C57BL/6J mice were orally inoculated with approximately 10^7^ cfu (n=4 per *Psychrobacter* accession, n = 8 for *P. lutiphocae*). Mice inoculated with the same strain were co-housed (Taconic) or single-housed (MPIDB) in sterile cages (IsoCage P, Tecniplast) and provided autoclaved water and sterile chow (NIH31M) ad libitum. Three weeks post-colonization, mice were euthanized and cecal contents were immediately collected, flash frozen, and stored at −80 °C. The Taconic experiments were performed in compliance with Taconic’s IACUC, and the MPIDB experiments were approved by and performed in accordance with the local animal welfare authority’s legal requirements.

To determine the bacterial colonization density in the mouse cecum, we serially diluted two aliquots per mouse of cecal material (50 mg each), incubated them on plates under their preferred conditions (Table S3) for three to five days. If no colonies were observed, we spread an inoculating loop of the undiluted aliquot onto Brain-Heart Infusion Agar and incubated at 37 °C for two weeks. For samples where we again observed no colonies, we categorized these *Psychrobacter* accessions as “non-persistent.” For the samples that did show colony growth, we confirmed the identity of the colonies as *Psychrobacter* using Sanger sequencing of 16S rRNA gene amplicons as described in the Supplemental Text.

### Statistical analysis

We performed all data processing and statistical analysis using R or Python. We compared means between groups using a Kruskal-Wallis test, followed by a pairwise Wilcoxon rank sum test to identify which groups differed when more than two groups were compared. We tested differences in frequencies between groups with more than 10 observations with a χ^2^-test, repeated a 100 times with down-sampling in order to correct for sampling sizes between groups. We measured phylogenetic signal using a log-likelihood ratio test on Pagel’s λ (Pagel’s λ fitted using the phylosig function from the phytools R package, and the null hypothesis being λ = 0). When applicable, we tested for the confounding of phylogeny with our groups of interest using the aov.phylo function from the R package geiger. We adjusted all p-values for multiple comparisons with the Benjamini-Hochberg (BH) correction method.

## Results

### *Psychrobacter* forms a clade whose basal members are of the *Moraxella* genus

To explore the evolutionary history of *Psychrobacter*, we built a phylogeny based on 400 conserved marker genes using publically available whole genomes derived from cultured isolates of 51 species from the Moraxellaceae family. We included 18 species each of *Acinetobacter* and *Moraxella* obtained from NCBI, and 15 *Psychrobacter* genomes that we generated in this study (Table S2). For all genomes, we incorporated phenotypic data collected from previously published type-strain research. Our analysis shows the 51 species formed three distinct clades with robust bootstrap support. The *Acinetobacter* clade consists uniquely of *Acinetobacter* species (labeled A in Fig 1A). The Acinetobacter clade is a sister taxon to the *Moraxella* (M) clade, consisting entirely of *Moraxella* species, and to the *Psychrobacter* (P) clade, which contains all of the *Psychrobacter* species, as well as four *Moraxella* species (*M. boevrei, M. atlantae, M. osloensis*, and *M. lincolnii*) that are basal in the P clade. This phylogeny indicates that these four *Moraxella* species are more closely related to *Psychrobacter* than to other *Moraxella*. P-clade *Moraxella* have more similar phenotypes compared to M-clade *Moraxella* than to *Psychrobacter* strains. Review of type strain descriptions revealed that *Psychrobacter* are consistently urease positive, nitrate reducing, salt tolerant, and non-fastidious, whereas P-clade *Moraxella* are inconsistent with their urease and nitrate reducing phenotypes, are sensitive to high salt concentrations, and are often nutritionally fastidious - they have complex growth requirements (in particular, often blood or bile for growth) [23, 24, 39, 42, 50, 51, 66, 67].

**Figure 1.**
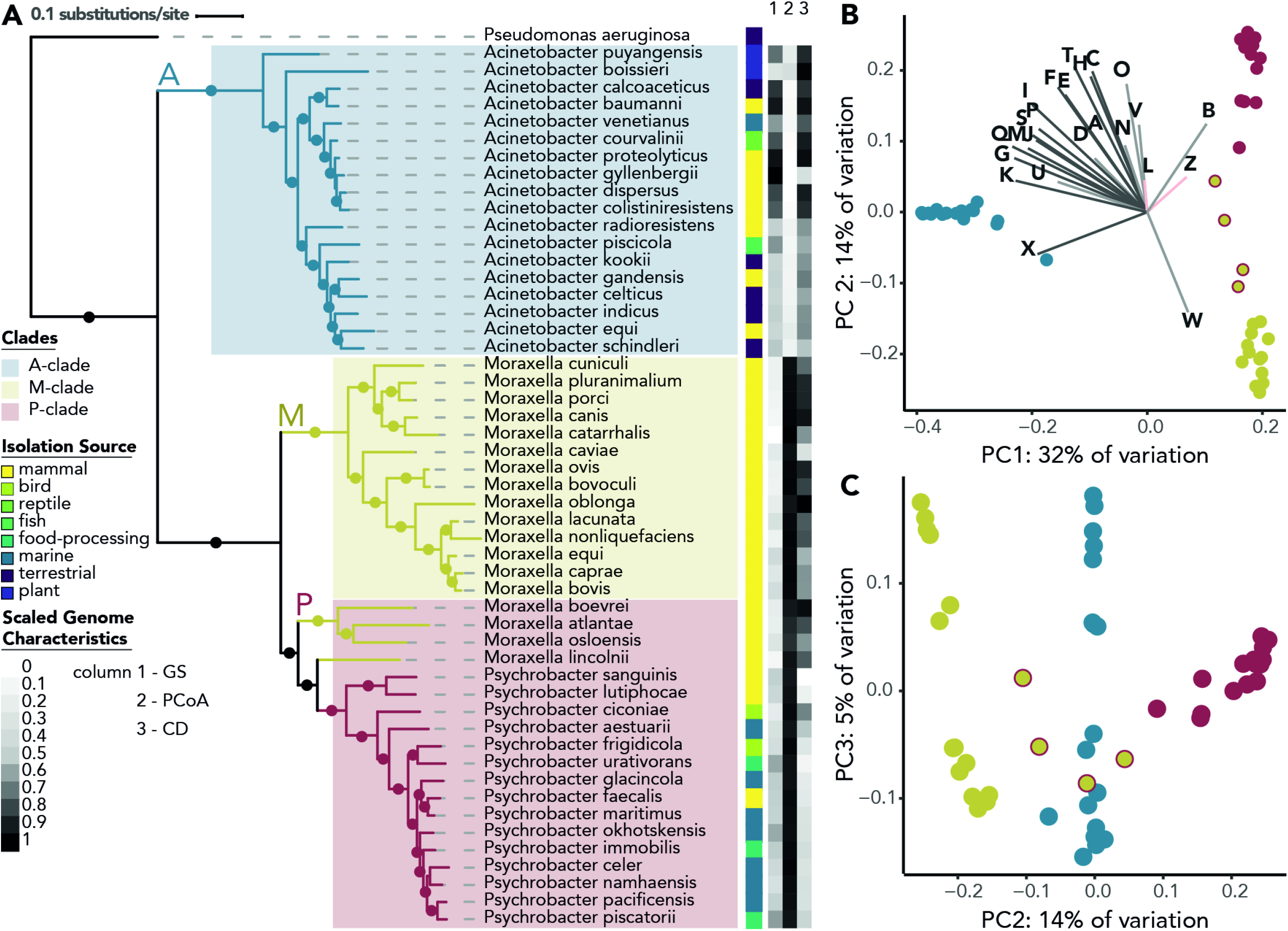
Genomic and phenotypic diversity in the family *Moraxellaceae*. A) The *Moraxellaceae* family phylogeny was constructed with 51 diverse *Moraxellaceae* genomes using the software PhyloPhlan, which constructs a phylogeny using fasttree with 1000 bootstraps, refined by RAxML under the PROTCATG model. Amino acid sequences from 400 marker genes were used in the alignment. Branches with bootstrap support greater than 70% are represented by filled circles. The scale bar represents the average amino acid substitutions per site. *Pseudomonas aeruginosa* was used as an outgroup. Clades are highlighted in colored blocks, and branches are colored by genus. Isolation source is depicted in a color strip, along with a heatmap of scaled notable genome characteristics that differ between the genera, with 0 representing the smallest value present and 1 the largest value (*P. aeruginosa* not included). GS = Genome size, ranging between 1.8 Mb and 4.5 Mb. PCoA = PC1 values from a PCoA based on gene presence/absence data. CD = genome coding density, ranging from 80% to 89%. Growth temperature range data was collected from type strain publications. B) PC1 and 2 of a PCoA analysis of a binary matrix of gene presence/absence for 51 species of *Moraxellaceae*, explaining 32% and 14% of the variation, respectively. Each genome is represented by one point, colored by genus. The P-clade *Moraxella* spp. are represented by yellow points with red outlines. The frequencies of genes associated with each COG category were associated with the PCoA axes as environmental vectors via the envfit function. All COG categories are significant (BH correction, p-value < 0.05), except L (replication, recombination and repair) and Z (cytoskeleton assembly and regulation) which are shown in pink vectors. COG categories with an r^2^ > 0.5 are shown in dark grey, while categories with an r^2^ < 0.5 are shown in light grey. C) PC2 and 3, explaining 14% and 5% of the variation.

Consistent with the topology of the phylogeny, a principal coordinates analysis (PCoA) of gene presence/absence data shows that the greatest variation within the family (*i.e*., Principle Coordinate [PC] 1) is the separation between the A clade versus the M and P clades, which are grouped (Fig. 1). The P-clade *Moraxella* spp. fall between the P-clade *Psychrobacter* and the M-clade *Moraxella* when visualizing PC1-2 (Fig 1B) as well as PC2-3 (Fig 1C). To assess what genes and gene functions may be contributing to the separation between the A-, M- and P-clades’ gene presence/absence, we performed an envfit function analysis using the R package vegan. This analysis indicates that the separation along PC1 and 2 is due to differences in shell genes (*i.e*., genes present in greater than two strains, but in fewer than 90%), not to differences in their core genes (*i.e*., genes present in between 90% and 100% of strains in the clade). Genes annotated with very diverse functions, falling under almost every Cluster of Orthologous Groups (COG) category, strongly contribute to the separation between clades (Fig. 1B, Table S5).

Despite their close phylogenetic relationship and high similarity in gene presence/absence, *Psychrobacter* and *Moraxella* have different genomic properties. When examined by genus rather than clade, *Psychrobacter* species have an average genome size of 3.12 ± 0.27 Mb, while the average of *Moraxella* is 2.41 ± 0.28 Mb (pairwise Wilcoxon rank sum test, p-value = 8e-07). *Moraxella* species have an average coding density of 86.4 ± 1.33%, which is significantly higher than the *Psychrobacter* species average of 82.7 ± 1.33% (pairwise Wilcoxon rank sum test, p-value = 1e-5). Despite being phylogenetically more related to *Psychrobacter* species than to the other *Moraxella* species, the P-clade *Moraxella* had significantly smaller genomes than *Psychrobacter* species and greater coding density (pairwise Wilcoxon rank sum test, p-values = 0.004) while not significantly different from the M-clade *Moraxella* spp. (pairwise Wilcoxon rank sum test, p-values = 0.9).

### *Psychrobacter* spp. ranges of growth temperatures differ from those of *Moraxella*

To examine the phenotypic behaviors of the *Moraxellaceae* family, we applied onto the previously generated phylogeny continuous trait mapping of the ranges of temperatures at which species from the *Moraxellaceae* family were reported to grow [23, 24, 27, 28, 35–75] (Fig 2). The *Psychrobacter* spp. included here are reported to have a broad range of growth temperatures (0 - 38 °C), but several strains, such as *P. frigidicola* and *P. glacincola*, are psychrophilic (restricted to growth below 20 °C), which is a phenotype that is not seen elsewhere in the family. Using the growth temperature information reported in the literature for this comparison, we observed that *Psychrobacter* spp. have lower minimum growth temperatures than *Moraxella* spp. from either the P- or M-clades (pairwise Wilcoxon rank sum test, p-value = 5e-06), which have a narrow range of temperatures at which they can grow (between 22 °C and 40 °C). In contrast to the minimum growth temperatures, there is little variation in the maximum growth temperatures, except for the notable exceptions of several *Psychrobacter* spp. that are restricted to growth at low temperatures.

**Figure 2.**
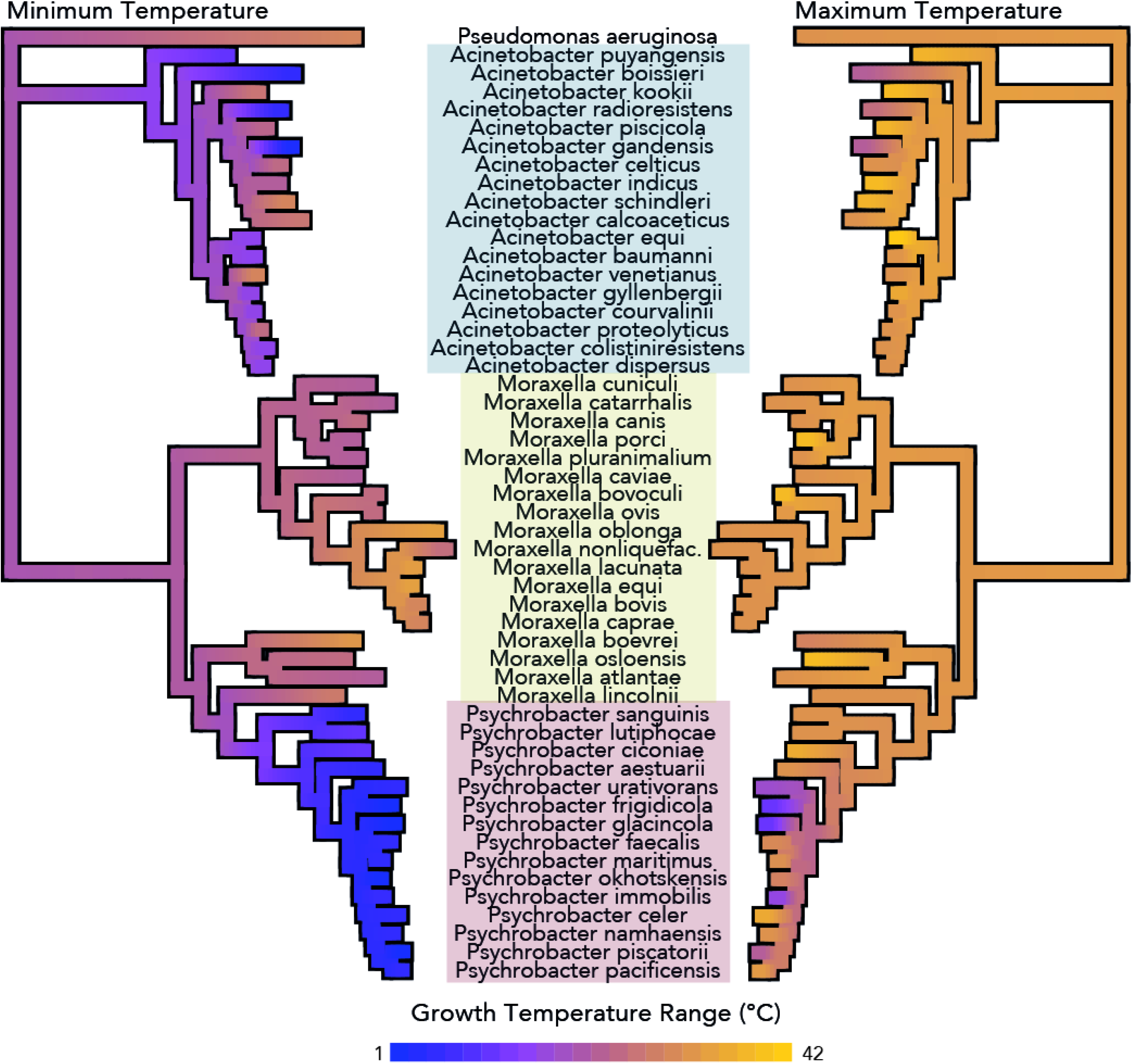
*Psychrobacter’s* restriction to cold temperatures is a newly emerged trait in the family *Moraxellaceae*. Continuous trait mapping growth temperature ranges of 51 species from the *Moraxellaceae* family, taken from type strain publications. Values at nodes are imputed by maximum likelihood analysis. The phylogeny was constructed by marker-gene analysis including 400 genes, as in Fig. 1. Genera are indicated with colored boxes.

### *Psychrobacter* spp. from diverse isolation sources show differences in cultivation temperatures

To explore further *Psychrobacte’s* phenotypic diversity, we established a strain collection of 85 *Psychrobacter* accessions isolated from diverse locations (Fig S1A) and ecological sources (Fig S1B). The optimal growth temperatures for each accession - provided by the catalogues from which the accessions were ordered - vary by isolation source (Kruskal-Wallis χ^2^ = 43.4, df = 9, p-value = 2e-06), with mammalian-derived strains reported as having significantly higher cultivation temperatures than strains from fish, invertebrates, sea water, terrestrial water, and soil samples (pairwise Wilcoxon rank sum test, adjusted p-values < 0.05) (Fig. S1C). Latitude of isolation has a confounding effect on reported optimal growth temperature, but explains very little of the variance (r^2^ = 0.06, F(1,67) = 5.5, p-value = 0.02). Examples of *Psychrobacter* morphology (P*. ciconiae* and *P. immobilis A351*) visualized by scanning and transmission electron microscopy are shown (Fig. S1D-G).

### Few *Psychrobacter* strains can grow at 37 °C

For a direct comparison of *Psychrobacter* phenotypes, we assessed the 85 *Psychrobacter* accessions for their ability to grow under 24 different combination of medium, salt concentration and temperatue (Methods & Supplemental Text). We calculated growth probabilities, or the fraction of growth positive conditions out of total conditions tested, for every strain and given variable of the growth curve screen, and compared them across the phylogeny and by isolation source. We generated a robust genus-level phylogeny for *Psychrobacter* using 400 conserved marker genes, with *M. lincolnii* as an outgroup (Fig. 3A). In agreement with single marker gene trees generated using *rpoB* sequences [34], 16S rRNA gene sequences [82], and the P-clade structure of *Moraxellaceae* family tree generated in this study, there is a phylogenetically basal group of strains mostly isolated from mammals, and a phylogenetically derived group isolated from mixed sources. Across the entire phylogeny, closely related strains have similar growth probabilities (Pagel’s λ ranging from 0.78 to 0.97, all corrected p-values < 1e-3). We observed that most *Psychrobacter* strains are tolerant of a wide variety of temperatures between 4 and 25 °C and of salt concentrations between 0 and 5%; more than 90% of all strains can grow under these conditions. However, only 54% of the tested accessions can grow at 10% added salt, and only 31% at 37 °C.

**Figure 3.**
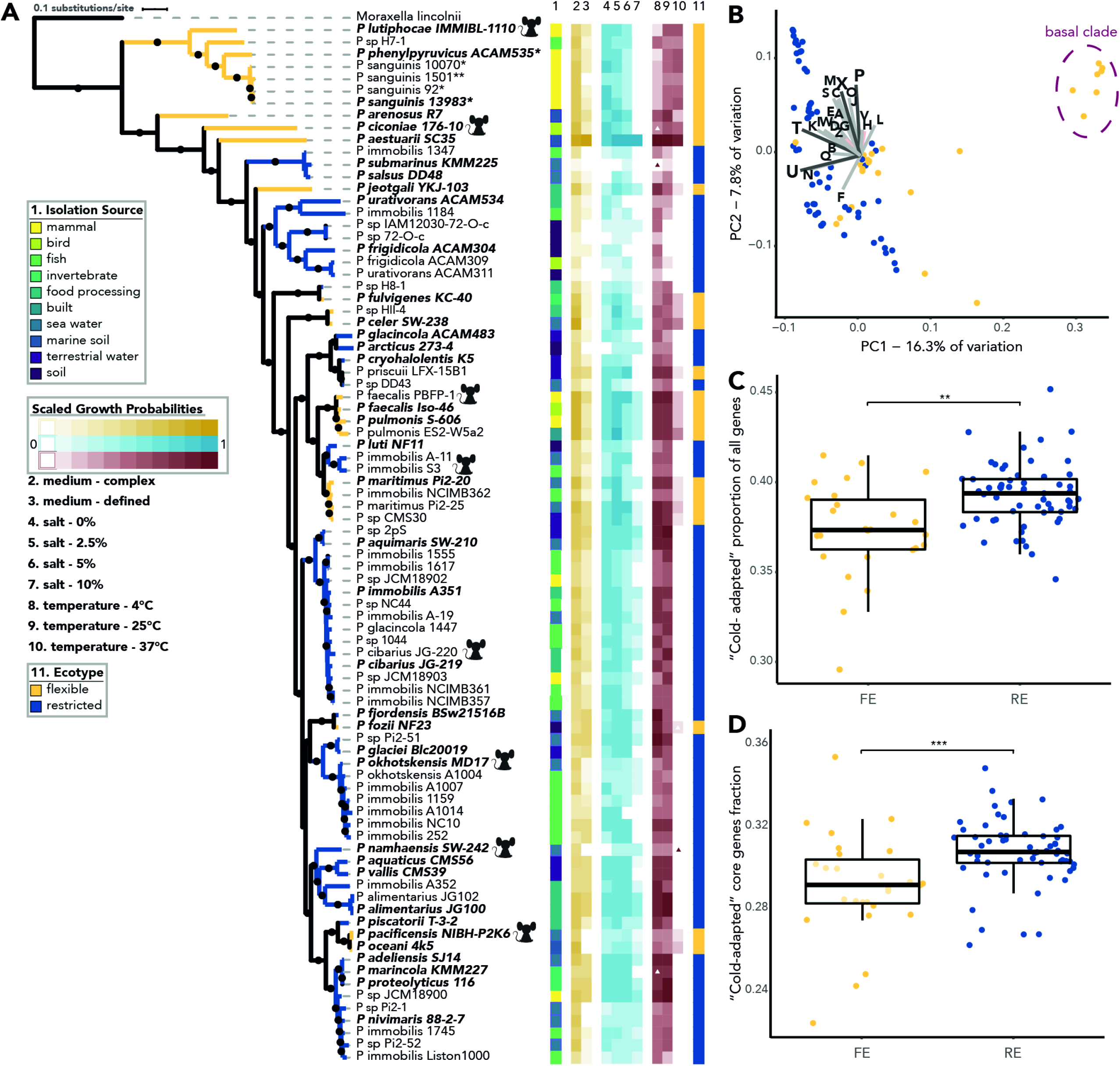
*Psychrobacter* phenotypic and genomic diversity. A) Using 85 *Psychrobacter* genomes, we constructed a genus-level phylogeny using fasttree with 1000 bootstraps, refined by RAxML under the PROTCATG model. Amino acid sequences from 400 marker genes were used in the alignment. Branches with bootstrap support greater than 70% are represented by filled circles. The scale bar represents the average amino acid substitutions per site. *M. lincolnii* was used as an outgroup. Type strain isolate names are indicated in bold and italicized type. Strains indicated with * next to their name exhibited growth defects in liquid media, and were tested on solid agar media instead. Strains indicated with ** exhibited growth defects on solid and liquid media, and were tested on solid media supplemented with 0.1% Tween80. Strains indicated with a mouse silhouette were later used in germ-free mouse colonization studies (Fig 5). Isolation source is depicted in column 1 as a color strip. Columns 2 – 10 represent the growth probabilities of each strain for each condition; media complexity is represented in yellow, salt concentration is represented in blue, and temperature is represented in red. Type strain data supports our temperature data except where indicated - colored triangles show conditions in which we expected growth but did not observe it, while white triangles represent conditions in which we observed growth we did not expect. The ecotype is shown in column 11 in a colorstrip, and in the color of the branches. B) The first two PCs of a PCoA of a gene presence-absence matrix of all 85 of the included accessions, colored by ecotype. The basal clade strains are shown within the ellipse. The frequencies of genes associated with each COG category were associated with the PCoA axes as environmental vectors. All COG categories are significant (BH correction, p-value < 0.05), except G (carbohydrate transport and metabolism) and H (coenzyme transport and metabolism) which are shown in pink vectors. COG categories with an r^2^ > 0.5 are shown in dark grey, while categories with an r^2^ < 0.5 are shown in light grey. C) The proportion of genes per genome falling in the highest quartile of “high cold adaptive amino acid traits” from each ecotype (n = 26 for FE, n = 59 for RE). D) the number of “core genes” (present in all 85 *Psychrobacter* accessions) that fall into the highest quartile of “high cold adaptive amino acid traits” (n = 26 for FE, n = 59 for RE). For all mean comparisons, the Wilcoxon rank sum test was used. ** indicates a p-value < 0.005, *** indicates a p-value < 0.0005.

Since 37 °C was the most restrictive condition tested, we divided the strains into two ecotypes: the “flexible ecotype” (FE) corresponds to strains that could grow at 37 °C, and the “restricted ecotype” (RE) corresponds to strains that could not grow at 37 °C. FE strains are psychrotrophic (mesophilic organisms with a low minimum growth temperature but an optimal growth temperature above 15 °C), and RE strains are either psychrotrophs or true psychrophiles (unable to grow at temperatures higher than 20 °C).

Notably, the basal clade of the *Psychrobacter-only* tree (Fig 3A) consists solely of FE strains, while the rest of the phylogeny is made up of a mixture of FE and RE strains. Furthermore, the basal FE strains have higher growth probabilities at 37 °C compared to other FE strains (Pagel’s λ = 0.89, p-value = 2e-5). Frequencies of RE and FE strains vary significantly across sources of isolation: the FE group is significantly enriched in strains derived from mammalian sources, and the RE group is significantly enriched in strains derived from fish, sea water and food sources (χ^2^-test, all p-values adjusted for group size < 0.05). Nonetheless, both FE and RE ecotypes contain strains from other environments, including mammalian-derived strains within the RE group. FE strains have higher growth probabilities at low-to mid-salt concentrations (Wilcoxon rank sum test, all p-values < 0.05), though there is no difference between FE and RE strains at 10% salt (Wilcoxon rank sum test, W = 811.5, p = 0.7). After accounting for phylogenetic relatedness, the growth probabilities under different salt concentrations are no longer significantly different between FE and RE strains (F(1,83) < 24.0, p-value < 0.4), indicating that ecotype grossly maps onto phylogeny. Finally, FE strains show higher growth probabilities in complex media compared to defined media, while RE strains showed little difference between the two (Wilcoxon rank sum test, W = 1.13e3, p-value = 0.0005). The difference in FE and RE strains’ growth probabilities under rich media remains significant after accounting for phylogenetic relatedness (F(1,83) = 43.4, p-value = 0.002).

### FE and RE *Psychrobacter* spp. have differences in genomic content

We looked for genes differentiating the FE and RE ecotypes by performing a microbial pan-genome wide association analysis (pan-GWAS) with the R package treeWAS [83] using gene presence/absence data. While this analysis returned no significant results, strong phylogenetic signals in the gene presence/absence data may have resulted in a loss of power for the pan-GWAS. When exploring gene presence/absence via PCoA, accessions from the basal FE-only subclade cluster closely together, indicating similar gene content. In agreement with the phylogeny, these basal FE accessions are separated from the other accessions on PC1, while the derived FE and RE accessions are more scattered, indicating more diverse gene content (Fig 3B). As with the separation in the *Moraxellaceae* PCoA of gene content, the separation between the basal FE-clade and the rest of the accessions is due largely to presence/absence patterns in shell genes (present in greater than one strain but fewer than 90%), not core (present in between 90 to 100% of strains) or cloud genes (present only in one strain), and all contributing genes were not unique to either clade. The separation is most strongly driven by genes from the COG categories T, signal transduction (r^2^ = 0.65, p-value = 0.001); U, trafficking and secretion (r^2^ = 0.63, p-value = 0.001); P, inorganic ion transport and metabolism (r^2^ = 0.56, p-value = 0.001), and X, unassigned or no homologs in the COG database (r^2^ = 0.51, p-value = 0.001)(Fig 5B, Table S5).

**Figure 4.**
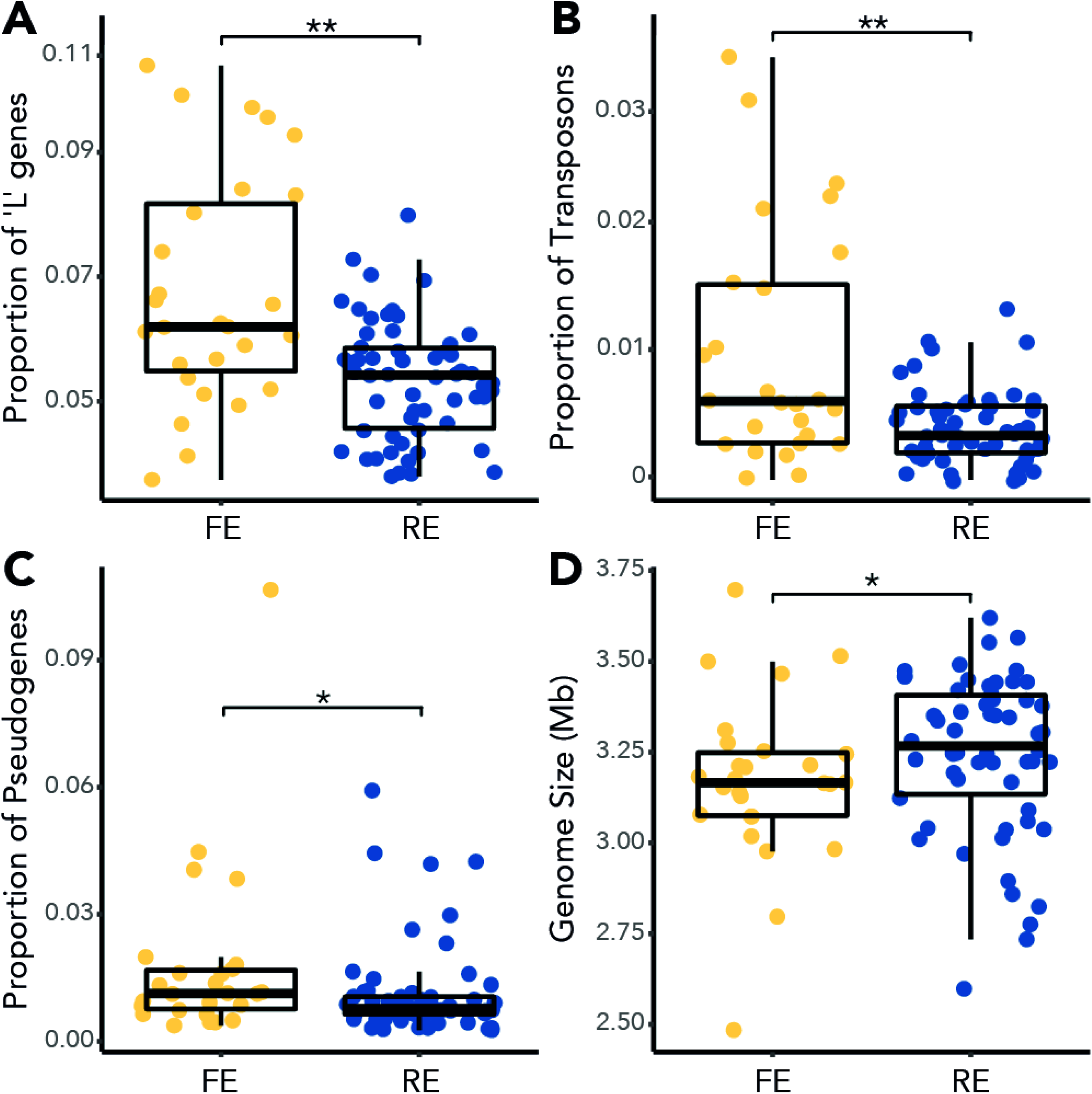
Divergence of ecotypes could be driven by transposon-mediated genome reduction. A) proportion of Cluster of Orthologous Groups (COG) category “L” (replication, recombination, and repair related) genes per genome. B) copy numbers per genome of all the transposases in the *Psychrobacter* pan-genome. C) Proportion of predicted pseudogenes per genome. D) *Psychrobacter* accession genome size in megabases. For all mean comparisons, the Wilcoxon rank sum test was used. * indicates a p-value < 0.05, ** indicates a p-value < 0.005. For each comparison, n = 26 for FE strains and n = 59 for RE strains.

**Figure 5.**
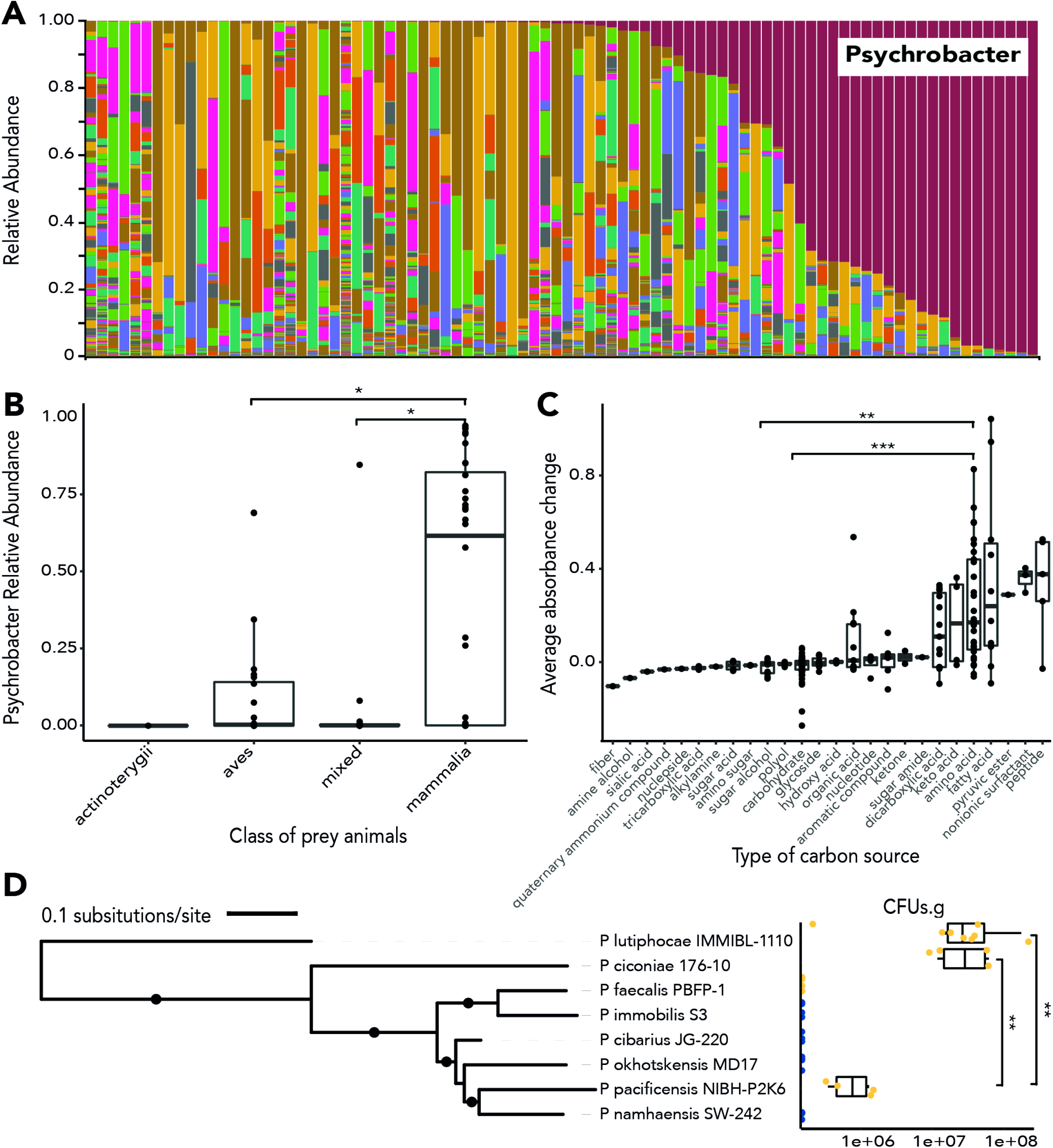
Psychrobacter strains occur and persist in the mammalian gut. A) Relative abundance at the genus level of 16S rRNA gene OTUs, clustered at 99% identity, of 86 polar bear fecal samples. *Psychrobacter* OTUs are colored red. B) *Psychrobacter* relative abundance in comparison to taxonomy of prey consumed by polar bears (n = 1 for *Actinoterygii*, n = 24 for *Aves*, n = 16 for mixed prey, and n = 32 for *Mammalia*). C) The average change in OD600 of 19 *Psychrobacter* accessions grown on 190 different substrates as sole carbon sources, compared across different classes of compounds. D) CFUs per gram of cecal contents of gnotobiotic mice is shown in comparison to accession phylogeny. 4 mice were tested per *Psychrobacter* strain, except *P. lutiphocae*, which was tested in 8 mice. FE strains are shown in yellow, while RE strains are shown in blue. The phylogeny was constructed as described in Fig. 3. Branches with bootstrap support higher than 70% are indicated with a filled circle. Means were compared using the Wilcoxon rank-sum test. *** indicates a p-value < 0.0005, ** indicates p-value < 0.005, * indicates p < 0.05.

We examined if there were qualitative differences in gene content between the FE and RE ecotypes, especially in functions related to host colonization versus cold adaptation. Both ecotypes carry genes associated with virulence [84–86] as well as genes that are potentially related to psychrophilic lifestyles [87] (Table S6).

We next tested whether there was a predicted difference in adaptation to cold environments in FE versus RE strains based on the amino acid properties of the protein-coding genes of their genomes. Amino acid traits have been implicated in psychrophilic lifestyles, including increased abundance of glycine, decreased abundance of proline, decreased arginine-to-lysine ratio, increased acidity, decreased isoelectric point, increased hydrophobicity, and increased GRAVY, when comparing homologs from psychrophiles to mesophiles or thermophiles [88–91]. Using these traits to define a protein sequence as cold-adaptive, REs have a higher proportion of cold-adapted proteins compared to FEs (Wilcoxon rank sum test, W = 407, p-value = 0.0006) (Fig. 3C). However, after including phylogenetic relatedness as a covariate, the difference is no longer significant (F(1,83) = 16.4, p = 0.2), indicating that cold adaptation in the RE is associated with their derivation from the basal strains. When comparing core proteins (present in 99 - 100% of accessions) RE strains have a higher percentage of cold-adapted core proteins than FE strains (Wilcoxon rank sum test, W = 404, p-value = 0.0004) (Fig. 3D), which is again confounded with phylogenetic relatedness (F(1,83) = 12.4, p = 0.2).

### FEs exhibit higher transposon copy numbers than RE strains

To determine whether there are genomic differences between the FE and RE ecotypes at a broader level than individual genes, we next examined the proportions of each genome devoted to each COG category. There are few differences between the ecotypes by COG category (Fig. S2). FEs have a significantly higher proportion of “L” category genes (replication-, recombination-, and repair-related genes) than REs (Wilcoxon rank sum test, W = 1.15e3, p-value = 0.0005; Fig. 4A). In particular, this difference stems from a higher proportion of transposon copies per FE genome than per RE genome (Wilcoxon rank sum test, W = 1.08e3, p-value = 0.003; Fig. 4B). As expected from the distribution of the RE and FE across the phylogeny, phylogenetic relatedness is confounded with ecotype in the effect on these difference (F(1,83) < 21.3, p = 0.1).

Increased transposon activity can lead to interruption and decay of functional protein-coding genes, leading to an increase in pseudogenes [92]. Given the higher number of transposons in FE strains, we next examined the number of predicted pseudogenes between the ecotypes. FE genomes are predicted to have a higher number of pseudogenes than RE strains (Wilcoxon rank sum test, W = 997, p-value = 0.03; Fig. 4C). Finally, we compared the average genome size between the ecotypes, as bacterial genomes are known to strongly select against the accumulation of pseudogenes [93]. FE strains have significantly smaller genomes than RE strains (Wilcoxon rank sum test, W = 524, p-value = 0.02; Fig. 4D). As with the other genomic properties, phylogeny confounds the comparison between the FE and RE groups’ pseudogene proportion and genome size (F(1,83) < 3.4, p-value < 0.6).

### Polar bear feces collected from the Arctic ice have high abundance of *Psychrobacter*

We surveyed the gut microbial diversity of 86 polar bear fecal samples, 76 wild and 10 captive, by 16S rRNA gene amplicon sequencing. *Psychrobacter* was detectable in 76/86 of the samples (Fig 5A). The large majority of *Psychrobacter* sequences (83%) were assigned to unclassified *Psychrobacter* spp.. We detected RE strain *P. immobilis* in 50% of samples with a mean abundance of 3%, and FE strain *P. pulmonis* in 8% of samples with a mean abundance of 0.5%.

Polar bear diet significantly impacted the abundance of *Psychrobacter* spp. (Kruskal-Wallis χ^2^ = 13.5, df = 3, p-value = 0.004); we found that polar bears feeding on mammalian prey, including seals and reindeer, had significantly higher abundances of unclassified *Psychrobacter spp*. than polar bears feeding on avian prey or mixed diets (pairwise Wilcoxon rank-sum test, adjusted p-values < 0.05) (Fig 5B). Unsurprisingly, diet data is confounded with location (Kruskal-Wallis χ^2^ = 8.6, df = 4, p-value = 0.0009) and year (Kruskal-Wallis χ^2^ = 16.6, df = 5, p-value = 0.005) of sample collection. Captive status did not significantly impact mean abundances, but there is a trend of wild bear samples having higher unclassified *Psychrobacter* spp. mean relative abundance than samples from captive bears (Wilcoxon rank sum test, W = 222, p-value = 0.08).

### Carbon source utilization patterns

To elucidate the effect that host dietary nutrition may have on *Psychrobacter* growth, we tested the maximum change in absorbance for 190 different carbon sources by a subset of *Psychrobacter* accessions including 9 FE and 10 RE strains (Fig 5C). All *Psychrobacter* spp. reached significantly higher OD_600_ growing on amino acid carbon sources compared to carbohydrates or sugar alcohols (pairwise Wilcoxon rank-sum test, adjusted p-values < 0.05). *Psychrobacter* spp. reach the highest OD600 growing on fatty acids, surfactants, and peptides. There was no significant difference between FE and RE strains’ changes in absorbance in these assays (Wilcoxon rank sum test, W = 1.55e6, p-value = 0.08).

### *Psychrobacter* strain survival in gnotobiotic mice

To assess the survivorship of *Psychrobacter* in a mammalian gut, we tested 8 accessions for persistence in the gastrointestinal tracts of germ-free mice (Fig. 5D). Chosen for phylogenetic breadth, we tested 4 FE strains, *P. ciconiae, P. faecalis PBFP-1, P. lutiphocae*, and *P. pacificensis*, and 4 RE strains, *P. cibarius JG-220, P. immobilis S3, P. namhaensis*, and *P. okhotskensis MD17*. Of the four FE strains tested, three were able to persist in the mice, while the FE strain *P. faecalis PBFP-1* and none of the RE strains were detectable after three weeks. Phylogenetic relatedness does not correlate with ability to colonize, as FE strain *P. pacificensis* was able to persist, while closely related RE strains, *P. namhaensis* and *P. okhotskensis*, were not. Phylogenetic placement does correlate with colonization density however, as the two most basal strains tested, *P. lutiphocae* and *P. ciconiae*, colonized at significantly higher densities than the most derived strain that was successful, *P. pacificensis* (pairwise Wilcoxon rank sum test, both adjusted p-values = 0.0007).

## Discussion

The phylogenomic and phenotypic characterizations of the *Psychrobacter* genus indicates a common ancestor with *Moraxella*, all of which are restricted to growth at higher temperatures. Furthermore, the most basal members of the *Psychrobacter* clade are *Moraxella* species and species of *Psychrobacter* that can grow at 37 °C, unlike most of the derived *Psychrobacter* species. Our extensive phenotyping indicated that members of the *Psychrobacter* genus grow at a wide range of salinities and temperatures, but it is the ability to grow at 37 °C that distinguishes strains the most, and which we used to define the two ecotypes, FE and RE. Our analysis of a large collection of wild polar bear feces shows both RE and FE strains are present, however tests in germfree mice support the notion that only FE may colonize the mammal gut, whereas RE may be allochthonous members or environmental contaminants. Together with previous reports, this work indicates the genus *Psychrobacter* is a lineage of pathobionts, some of which have evolved to inhabit the colder environments of their warm-bodied hosts.

Our results corroborate those of Bakermann, who used the isolation source of *Psychrobacter* as a proxy for temperature adaptation to conclude the genus has a mesophilic ancestor [34]. By assessing growth under the same 24 conditions for 85 strains, we remove any ambiguity that can stem from whether an isolate can indeed grow at the temperature of its source of isolation. This is particularly important in light of our results showing that several strains isolated from mammals proved to be RE, and that many of the FE strains came from sea water or other relatively cold environments.

*Psychrobacter’s* sister taxon *Moraxella* in particular is commonly isolated from host mucosal tissues, and exhibits the reduced genome size and nutritional fastidiousness common to many host-dependent organisms. *Moraxella* contains species that are frequently associated with human respiratory infections, primarily *M. catarrhalis* [84], as well as livestock conjunctivitis, for example, *M. bovis* or *M. equis* [94]. Since they are commonly found in healthy individuals and can cause disease in healthy individuals [95], *Moraxella* are best categorized as pathobionts and not dedicated or opportunistic pathogens. Several species of *Moraxella* appear basally in the P-clade of the *Moraxellaceae* family-level phylogeny, suggesting that *Psychrobacter* evolved from a “*Moraxella*-like” ancestor. This is supported by the fact that both phylogenetically basal and derived *Psychrobacter* strains carry genes related to virulence functions, and that many of the basal *Psychrobacter* strains exhibit growth defects in liquid culture, similar to the fastidiousness of *Moraxella*.

Despite clear phenotypic differentiation, *Psychrobacter* and *Moraxella* have similar genomic content, although *Psychrobacter* genomes are larger. A psychrophile emerging from an apparently mesophilic background through widespread horizontal gene transfer has been suggested before in the genus *Psychroflexus* [96], though the study was limited to comparing two genomes. In fact, it has been suggested before that this is *Psychrobacter’s* evolutionary trajectory [34], and although many *Psychrobacter* trees are constructed using a *Moraxella* outgroup, *Psychrobacter’s* potential pathogenic origin has not been widely discussed. Horizontal gene transfer would explain *Psychrobacter’s* larger genome size compared to *Moraxella* despite lower coding density, as many newly acquired horizontally transferred genes are expected to be inactivated and pruned by the recipient genome [96, 97].

Our data show that the largest phenotypic divide within the genus is the ability to grow at 37 °C, which we used to sort strains into FE and RE. FE strains make up the basal clade of the *Psychrobacter* phylogeny. Their smaller genomes show less cold-adaptation in their protein-coding genes than RE strains with proportionally fewer cold-adaptive proteins and more transposons. The three of four FE strains tested were able to colonize germ-free mice, whereas none of the RE strains could, indicating that growth at 37 °C may be necessary (although not sufficient) to colonize mammals. Opportunistic infections in mammals caused by *Psychrobacter* strains are limited to *P. sanguinis, P. phenylpyruvicus, P. faecalis*, and *P. pulmonis* [32], which while are all FE strains. Our results suggest that the FE strains are maintaining an ancestral ability to grow at mammalian body temperatures and colonize mammalian host bodies, while RE strains have adopted a psychrophilic lifestyle.

Adaptation to psychrophilic lifestyles usually arises conjointly with high tolerance for salt, but at very high salt concentrations, RE strains had similar growth probabilities as FE strains. Both FE and RE strains carry genes relating to both salt tolerance and cold adaptation functions, such as compatible solute accumulation or membrane fluidity control. The similar salt tolerances of FE and RE strains is thus not surprising, given that both ecotypes have enough psychrophilic adaptation to grow well at 4 °C. There may be a difference between ecotypes in salt tolerance and cold tolerance in more extreme conditions than those we tested.

Phylogenetically basal FE strains show stronger growth at 37 °C than phylogenetically derived FE strains. It may be that genes responsible for ancestral strains’ ability to grow at 37 °C were lost between the differentiation of the basal and derived clades. Subsequently derived *Psychrobacter* strains could have then acquired other genes providing the same FE phenotype as the basal strains, making pinpointing the genes responsible for the FE phenotype difficult. Genes that are unannotated and do not have homologs in databases such as UniRef or COG may also play a role in phenotypic and genomic differentiation of FE and RE strains, so future *in vitro* phenotypic screens may be helpful in assigning function to the many hypothetical genes in the *Psychrobacter* pan-genome.

The difference in ecotypes could also be due to gene regulation rather than gene presence and absence. There has been some transcriptional work done in *P. arcticus* [98], but it focused on comparing gene expression at ambient temperatures (around 20 °C) to ultra-low temperatures (−10 °C) to study *P. arcticu*s’s evolutionary approach to cold-adaptation. Future studies might use the same techniques to identify differences in gene expression responsible for FE strains’ phenotypic plasticity by comparing gene expression at low temperatures (for example, 4 °C) to higher temperatures (37 °C).

Our data reveal clear genomic signatures between the ecotypes. In particular, we observed higher transposon proportions and smaller genomes in FE strains compared to RE strains. Transposon-mediated genome reduction in host-associated bacteria compared to their free-living relatives has been observed in as-of-yet unculturable light-producing symbionts from ceratoid deep-sea anglerfish [99]. While none of the *Psychrobacter* spp. exhibit the fastidiousness of truly host-dependent bacteria [100], it is striking that FE strains were more likely to grow in nutritionally complex media than defined media, compared to the RE strains, for which nutritional complexity had little impact.

It is possible that FE strains have enough contact with mammalian hosts that it remains advantageous for them to maintain their ability to grow at higher temperatures in rich nutritional environments. *Psychrobacter* spp. have been reported previously in the skin, respiratory, and gut microbiomes of several marine mammals, including whales, porpoises, seals, and sea lions; it could be argued that *Psychrobacter* presence is due to contamination from sea water. Polar bears hunt seals on sea ice and their diet during the summer includes bird eggs. Our data show that polar bears consuming seal meat have higher *Psychrobacter* abundance than those consuming eggs, which may result from their time on the sea ice, particularly since FE and RE were equally represented. *Psychrobacter* spp. grew to high densities when grown on amino acids, peptide mixtures, and fatty acids: all carbon sources that would be abundant in the gut of a polar bear eating fatty seal meat. This may allow FE *Psychrobacter* strains to thrive in the bear gut. Given that the majority of the *Psychrobacter* diversity detected from the polar bear feces could not be classified, much remains to be learned about the natural history of this genus.

The history of the genus *Psychrobacter* is one of an ancestral pathobiont or pathogen, some of the descendants of which attenuated their own pathogenicity to broaden their ecological distribution. The emergence of a psychrotroph - a remarkable generalist - from a background of a more specialized pathobiont or pathogen showcases the adaptability of bacteria, and particularly *Proteobacteria*, to their environments.

## Data availability

Raw sequences for the *Psychrobacter* genome sequencing and polar bear feces 16S rRNA gene sequencing, as well as assembled *Psychrobacter* genomes, are available in the European Nucleotide Archive under the accession PRJEB40380. Annotated *Psychrobacter* genomes are available at ftp://ftp.tue.mpg.de/pub/ebio/dwelter. Raw data, R notebooks, and Python scripts for the analyses are available at https://github.com/dkwelter/Welter_et_al_2020.

## Acknowledgments

This work was supported by the Max Planck Society. We thank Jacobo de la Cuesta-Zuluaga, Sara Di Rienzi, Hagay Enav, Angela Poole, Jessica Sutter, Taichi Suzuki, and William Walters for discussions regarding project design and analysis, and Andrea Belkacemi, Ilja Bezrukov, Pablo Carbonell, Silke Dauser, Julia Hildebrandt, and Christa Lanz for their advice on and assistance with sequencing. We also thank Jürgen Berger and Katharina Hipp for performing electron microscopy. We thank Markus Dyck and Patricia Morin for providing the Polar Bear Habitat samples, Maria Frank for samples from the Metro Toronto Zoo, Daryll Hedman and Manitoba Conservation for the sample from the Polar Bear Holding Facility in Churchill, and Sam Iverson for samples from Hudson Straight. The collection of Nunavut samples would not have been possible without collaboration of colleagues at the Gjoa Haven Hunters and Trappers Association and their Traditional Ecological Knowledge relating to polar bears. The Nunavut field work was supported by funds from the Nunavut Wildlife Management Board (NWMB), the Nunavut General Monitoring Plan (NGMP), the National Science and Engineering Research Council (NSERC), and Environment Canada (Gov. of Canada). We would also like to thank Marie Pages and Maxime Galan for assistance with cytochrome b barcode sequencing.

