## Supplementary material for "Free-living psychrophilic bacteria of the genus *Psychrobacter* are descendants of pathobionts": Table S2

Table S1 - Software versions and parameters used in analyses

| **Purpose** | **Software** | **Version** | **Parameters, if not default** |
| --- | --- | --- | --- |
| **Moraxellaceae family genomics and phenotypic data** | PhyloPhlAn [1] | v2.0 | diversity = ‘medium’, and set to ‘accurate’ |
|  | CheckM [2] | v1.0.18 |  |
|  | Prokka [3] | v1.14.6 | kingdom = ‘bacteria’ |
|  | interactive Tree of Life (iTOL) [4] | v5.6.1 |  |
|  | PanX [5] | v1.6.0 | core gene cutoff = 90% |
|  | MCL | v14.137 |  |
|  | eggNOG mapper [6] | v1.0.3 |  |
| **Genome sequencing, assembly and annotation.** | fqtools | v2.0 |  |
|  | clumpify | v37.78 | dedupe = t, dupedist = 40 for HiSeq, 2500 for MiSeq, optical = t |
|  | bbduk | v37.78 | minimum read length = 100 bp, minimum PHRED quality score = 25 |
|  | skewer | v0.2.2 |  |
|  | fastqc | v0.11.7 |  |
|  | multiqc | v1.7 |  |
|  | seqtk | v1.3 | number of sub-sampled reads per sample = 1 000 000 |
|  | bbnorm | v37.78 | target = 50, k = 31, minkmers = 15, prefilter = t |
|  | SPAdes | v3.12.8 | cov_cutoff = off, set to careful, minimum scaffold length = 500 bp |
|  | pilon | v1.22 | chunksize = 10 000 000 |
|  | quast | v5.0.0 |  |
|  | sourmash | 2.0.0a4 | scaled = 10 000, k = 31 |
|  | gtdbtk | v1.0.2 | min_perc_aa = 10 |
|  | MinKNOW | v3.5.5 |  |
|  | ont-guppy | v3.2.4 |  |
|  | porechop | v0.2.4 | adapter_threshold = 90, minimum PHRED quality score = 8, minimum read length = 500 bp |
|  | NanoPack - nanocomp | v1.11.3 |  |
|  | NanoPack - nanofilt | v2.7.1 |  |
|  | NanoPack - nanoget | v1.14.0 |  |
|  | NanoPack - nanolyse | v1.1.3 |  |
|  | NanoPack - nanomath | v0.23.3 |  |
|  | NanoPack - nanoplot | v1.31.0 |  |
|  | NanoPack - nanostat | v1.2.1 |  |
|  | Unicycler | v0.4.8 | minimum contig length = 500 bp, mode = normal |
|  | DFAST [7] | v1.2.6 |  |
|  | DIAMOND [8] | v0.9.24 |  |
|  | UniRef90 [9] | downloaded April 2018 |  |
| **Microbiome diversity of polar bear feces.** | QIIME [10] | v2 | DADA2 quality control (F read trimmed to 200 bp, R read trimmed to 110 bp), samples rarefied to 20 000 sequences |
|  | Silva [11] | 138 SSURef NR99 515F/806R | |
| **Statistical analysis.** | Python | v3.6.10 |  |
|  | Biopython - Protein Analysis class [12] |  |  |
|  | R [13] | v3.6.3 |  |
|  | R package ecodist [14] | v0.3.0 |  |
|  | R package geiger [15] | v2.0.7 |  |
|  | R package phytools [16] | v0.7-47 |  |
|  | R package treeWAS [17] | v1.0 |  |
|  | R package vegan [17] | v2.5-6 |  |
