## Supplement for "Free-living psychrophilic bacteria of the genus *Psychrobacter* are descendants of pathobionts"

**Supplemental Material**

**Materials and Methods**

**Electron microscopy.** We picked two strains of *Psychrobacter* (*P. ciconiae* and *P. immobilis* strain *A351*) for visualization. We inoculated three biological replicates in a defined medium used for *Psychrobacter* [1], grew the cultures to mid-log phase at 25 ^o^C, spun down the cells, then washed them twice in PBS. Cultures were fixed and imaged by the Electron Microscopy Center at the Max Planck Institute for Developmental Biology. Cells were fixed with a mixture of 4% formaldehyde and 2.5% gluteraldehyde. For scanning electron microscopy, cells were postfixed with 1% osmium tetroxide on ice for one hour, then dehydrated in a graduated series of ethanol followed by CO_2_ critical point drying with the Polaron E3000 (Quorum Technologies, Lewes, United Kingdom). Cells were sputter-coated with a 6 nm thick layer of platinum and examined with the S-800 field emission scanning electron microscope (Hitachi High Technologies, Tokyo, Japan) at an accelerating voltage of 15 kV. For transmission electron microscopy, cells were embedded in low melting agarose and sectioned into 1 mm^3^ cubes. Cells were postfixed first with 1% osmium tetroxide on ice for one hour, followed by 1% uranyl acetate on ice for one hour, then dehydrated in ethanol and embedded in epon resin. Ultra-thin sections were stained with uranylacetate and lead citrate, and examined with a Tecnai Spirit transmission electron microscope (Thermo Fisher Scientific) operated at 120 kV.

***Psychrobacter* phenotypic screen.** We used Lysogeny broth (LB) as a complex medium with high nutrient availability, and a variation of M9 minimal medium (MM) as a defined media with lower nutrient availability. We prepared LB following manufacturer instructions except by varying concentrations of sodium chloride (NaCl) (see below), and autoclaving it at 121 °C for 20 minutes to ensure sterility. We prepared MM by adding components in the following final concentrations, followed by filter sterilization: 33.7 mM Na_2_HPO_4_, 22.0 mM KH_2_PO_4_, 9.35 mM NH_4_Cl, 0.4% L-glutamic acid, 1 mM MgSO_4_, 0.3 mM CaCl_2_, 1x ACTC trace vitamins solution, and 1x ACTC trace minerals solution, pH 7 (adjusted with KOH). We added NaCl to each ‘base medium’ to one of the four following concentrations of NaCl - low, with 0%, medium, with 2.5%, medium-high with 5%, and high with 10%. We grew cultures at 4 ^o^C, representing cold environments, 25 ^o^C representing mesophilic environments, and 37 ^o^C representing mammalian host body temperature.

*P. phenylpyruvicus* and *P. sanguinis* exhibited the inability to grow in liquid culture under the conditions tested. Hence, we streaked five strains of these two species first on Columbian Blood Agar which we incubated for 3 days at 37 ^o^C, then we washed and diluted them as for the liquid cultures. We spotted dilutions onto LB or MM media with 3% agar, supplemented with the tested salt concentrations as above, and incubated at 4 ^o^C, 25 ^o^C, and 37 ^o^C. *P. sanguinis* strain *1501* was unable to grow on any of the base media, so we repeated the same process with media supplemented with 0.1% Tween80. We scored growth for all plates after 14 days.

For two accessions, we could not confirm the purity of the cultures used in the phenotypic screen, and subsequently removed them from analysis. They were removed from subsequent genomic analysis as well.

For the type strains included in the phenotypic screen, we cross-checked growth probabilities at 4 ^o^C and 37 ^o^C with the published type strain descriptions where the original publications were clear about conditions tested. For the majority of strains, the type strain data and our data are in agreement; however, there are discrepancies between the type strain publications and our own data for the following strains: *P. ciconiae*, *P. fozii*, *P. marincola*, *P. namhaensis*, and *P. submarinus*. In some cases, we observed growth under temperatures where growth was previously unobserved; *P. ciconiae* and *P. marincola* have not before been reported to grow at 4 ^o^C, and *P. fozii* has not been reported to grow at 37 ^o^C. In other cases, we observed no growth in temperatures that have previously been reported to support growth; *P. submarinus* is expected to grow at 4 ^o^C and *P. namhaensis* is expected to grow at 37 ^o^C. The conditions tested in the type strain publications often differ from the conditions used in our phenotypic screen.

***Psychrobacter* carbon utilization assay.** To inform the design of the minimal medium used in the phenotypic screen described above, we chose a subset of 19 *Psychrobacter* strains (*P. adeliensis, P. aestuarii, P. aquaticus, P. arenosus, P. celer, P. cibarius JG-219, P. ciconiae, P. cryohalolentis, P. faecalis PBFP-1, P. fozii, P. fulvigenes, P. glacincola ACAM483, P. luti, P. lutiphocae, P. maritimus Pi2-25, P. okhotskensis MD17, P. pacificensis, P. piscatorii, P. urativorans ACAM534*) to evaluate for their ability to grow on 190 different substrates as their sole carbon source. We utilized Biolog plates PM1 and PM2 (Biolog Incorporated, Hayward, CA, USA) following a slightly modified protocol.

Briefly, we streaked out strains on agar plates of their preferred medium incubated at their preferred temperature (Table S3). We scraped cells from these plates and resuspended them at a final optical density at 600 nm (OD_600_) of 0.07 in the Biolog inoculation fluid (PM IF-0a GN/GP 1.2x) diluted to 1x with sterile water. We inoculated 100 uL of cell suspension into each well of the PM1 and PM2 plates, mixing well to resuspend the carbon sources. All strains were grown under aerobic conditions at their preferred growth temperature (Table S3). Growth was monitored by measuring OD_600_ for 14 days. We calculated total change in absorbance by subtracting the blank wells from the substrate wells, then taking the difference between the absorbances at t = 14 days and t = 0 (inoculation time point). We then averaged the total change in absorbance for all strains for every carbon source, and assigned each carbon source to a “family” of compounds.

All strains reached a significantly higher max OD_600_ in amino acid substrates compared to other carbon sources. L-glutamate was chosen as the carbon source for the defined medium, as none of the strains tested failed to grow using it. Some of the other amino acids allowed strains to grow to a larger change in OD_600_, but failed to allow all strains to grow. The minimal medium we used in our phenotypic screen was designed based off of the M9 minimal medium, replacing glucose with glutamic acid, and varying salt concentrations as described above.

**Isolation of *Psychrobacter* sp. from polar bear feces.** Two wild polar bear fecal samples were combined and diluted to 1 mg/mL in PBS. We plated the solution onto LB agar + 6% NaCl and incubated it at 14°C for 10 days. A single colony grew and was identified as *Psychrobacter faecalis* by colony PCR and Sanger sequencing of the full length 16S rRNA gene (described below). This isolate is designated as *P. faecalis PBFP-1* and is included in the phenotypic screen described above and the genomic analysis.

**Genome sequencing, assembly and annotation.** We initially sequenced samples using the MiSeq 2x250 bp and the HiSeq 2x150 bp paired-end read technology (Illumina, San Diego, CA, USA) as previously described [2] (Table S4). We constructed libraries using the Nextera DNA Sample Preparation Kit (Illumina) with modifications: we sheared 1 ng of DNA with in-house-generated Tn5 transposase, then amplified and barcoded it with custom primers for 7 to 14 cycles. We pooled samples and size-selected using magnetic beads for MiSeq libraries or BluePippin (Sage Science, Beverly, MA, USA) for HiSeq libraries. After dilution to 4 nM for MiSeq and 2.5 nM for HiSeq, we stored libraries at -20 °C until sequencing.

After sequencing and demultiplexing, we validated raw reads with fqtools, and de-duplicated them with Clumpify. bbduk and Skewer were used to remove sequencing adapters and filter reads. At multiple steps throughout, we used fastqc and multiqc to monitor quality. After quality control, reads were assembled *de novo*. First, we subsampled reads using seqtk, normalized by bbnorm, and then assembled with SPAdes, followed by refinement with Pilon. We ran MultiQC and QUAST after assembly and refinement. Finally, we assessed the assemblies for quality using CheckM. We assigned taxonomy using Sourmash and GTDB-Tk, and annotated assemblies using Prokka.

For genomes that were particularly fragmented (having greater than 250 contigs), we performed additional long-read sequencing (Supplementary Table 4). We constructed Oxford Nanopore libraries using the Ligation Sequencing and Native Barcode Ligation kits (Oxford Nanopore, Oxford, UK). We sequenced the libraries using the MinION® system run with software MinKNOW (Oxford Nanopore). We basecalled and demultiplexed reads using ont-Guppy, and used Porechop [3] to ensure that the adapters were removed, along with poor quality reads. We generated summary statistics, summary plots, and removed lambda phage reads using NanoPack [4]. After quality control, we combined the long reads with the short (generated by the MiSeq and HiSeq libraries described above) for hybrid assembly. We assembled and analyzed the hybrid assemblies in largely the same way as the short-read assemblies described above, however replacing SPAdes with Unicycler [5].

We followed the quality control cutoffs suggested by CheckM, and removed two genomes for having contamination higher than 5%, and one for having completion less than 90%. We removed two additional genomes as the taxonomic classification was outside of the *Psychrobacter* genus. These accessions were removed from genomic and phenotypic analysis. For a full description of sequencing for each accession, see Table S4.

**16S rRNA gene Sanger sequencing.** To confirm the identity of isolates as *Psychrobacter*, we performed full-length 16S rRNA gene sequencing. We prepared “colony” polymerase chain reactions following the protocol for Phusion High Fidelity Polymerase (New England Biolabs, Ipswitch, MA, USA), swirling a colony of interest in the reaction mixture as a substitute for the DNA template, and the 27F and 1391R universal full length 16S rRNA gene primers [6] for amplification. The reactions were incubated on the Mastercycler pro S thermocycler (Eppendorf, Hamburg, Germany) following the Phusion protocol, using a touchdown program for the annealing temperature (dropping the annealing temperature at a rate of 1 ^o^C per cycle from 70 ^o^C to 55 ^o^C, then annealing at 55 ^o^C for 15 cycles). We cleaned the products using the DNA Clean & Concentrator -25 kit (Zymo Research, Irvine, CA, USA) and checked the concentration using the DS 11+ Spectrophotometer (DeNovix, Wilmington, DE, USA). We used the cleaned DNA products as the templates for the Sanger sequencing reaction, following the ABI PRISM BigDye Terminator Cycle Sequencing Kit (ThermoFisher Scientific, Waltham, MA, USA) protocol with the 27F primer. The sequencing was performed using the 3730xl DNA Analyzer (ThermoFisher). Upon sequencing completion, we BLASTed [7] the sample sequences against the NCBI non-redundant nucleotide database. We examined the top ten hits to confirm the sample identity.

1.	Bergholz PW, Bakermans C, Tiedje JM. Psychrobacter arcticus 273-4 uses resource efficiency and molecular motion adaptations for subzero temperature growth. *J Bacteriol* 2009; **191**: 2340–2352.

2.	Karasov TL, Almario J, Friedemann C, Ding W, Giolai M, Heavens D, et al. Arabidopsis thaliana and Pseudomonas Pathogens exhibit stable sssociations over evolutionary timescales. *Cell Host Microbe* 2018; **24**: 168–179.e4.

3.	Wick R. Porechop. Github.

4.	De Coster W, D’Hert S, Schultz DT, Cruts M, Van Broeckhoven C. NanoPack: visualizing and processing long-read sequencing data. *Bioinformatics* 2018; **34**: 2666–2669.

5.	Wick RR, Judd LM, Gorrie CL, Holt KE. Unicycler: Resolving bacterial genome assemblies from short and long sequencing reads. *PLoS Comput Biol* 2017; **13**: e1005595.

6.	16S ribosomal DNA | Lutzoni Lab. <http://lutzonilab.org/16s-ribosomal-dna/.> Accessed 13 Jan 2020.

7.	Altschul SF, Gish W, Miller W, Myers EW, Lipman DJ. Basic local alignment search tool. *J Mol Biol* 1990; **215**: 403–410.

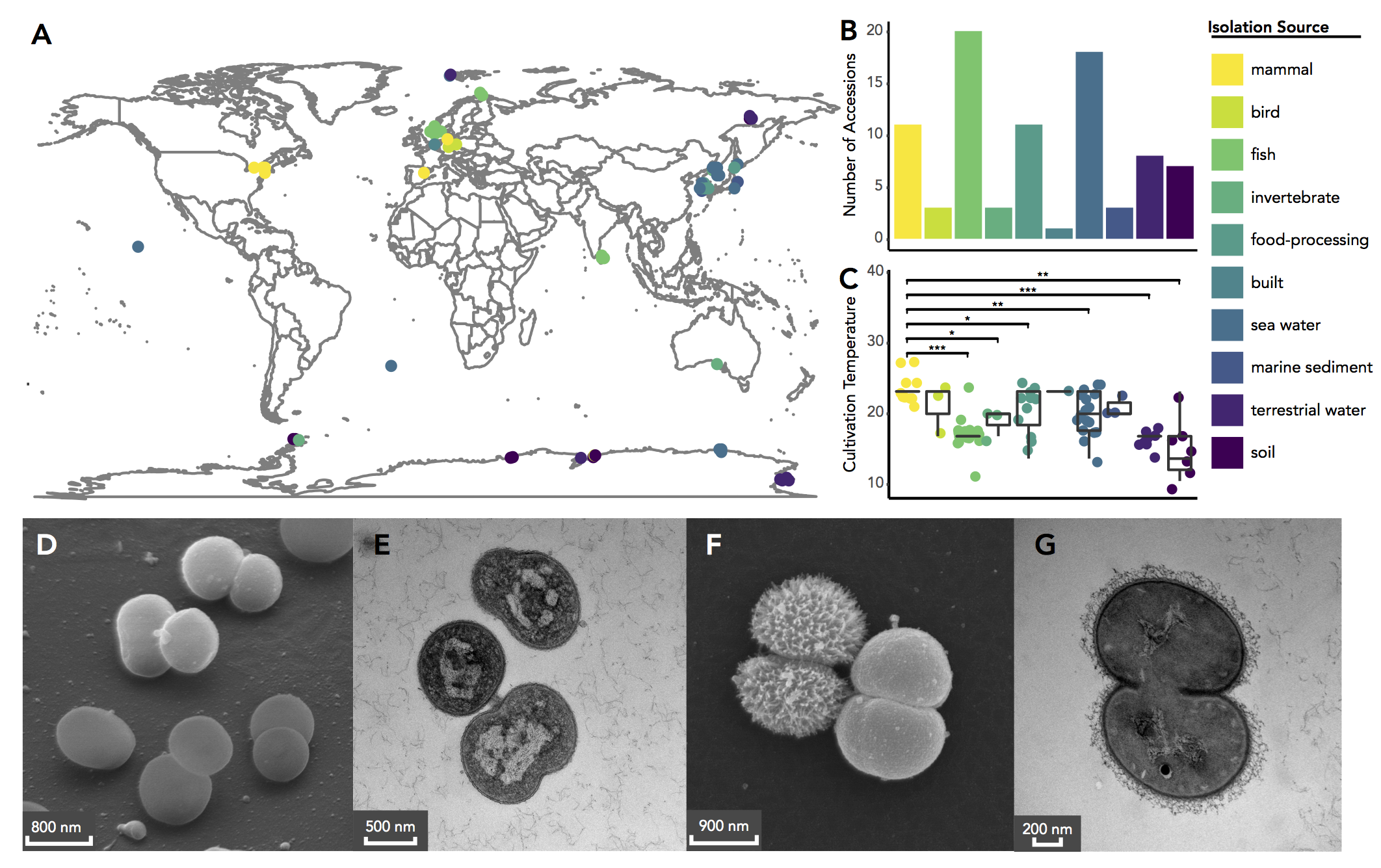

**Figure S1. Psychrobacter spp. can be isolated from diverse sources and many locations.** A) geographic locations of isolation of the strains used in this study, provided by the strain catalogue the accessions were purchased from when available. B) count of accessions from each of the following isolation sources: mammal, bird, fish, and invertebrate host bodies, processed food stuffs (including frozen and fermented foods), the built environment, terrestrial sources including freshwater and soil, and marine sources including sea water or ice, and marine sediment. C) cultivation temperatures from each isolation category, as provided by the strain catalogues. Means were compared using the pairwise Wilcoxon rank sum test. * indicates p < 0.05, ** p < 0.005, and *** p < 0.0005. D and F)  Scanning electron microscopy of P. immobilis strain A351 and P. ciconiae grown at 25 ^o^C in a defined medium. E and G) Transmission electron microscopy of P. immobilis and P. ciconiae under the same conditions. Images are representative of biological triplicates and multiple fixation techniques.

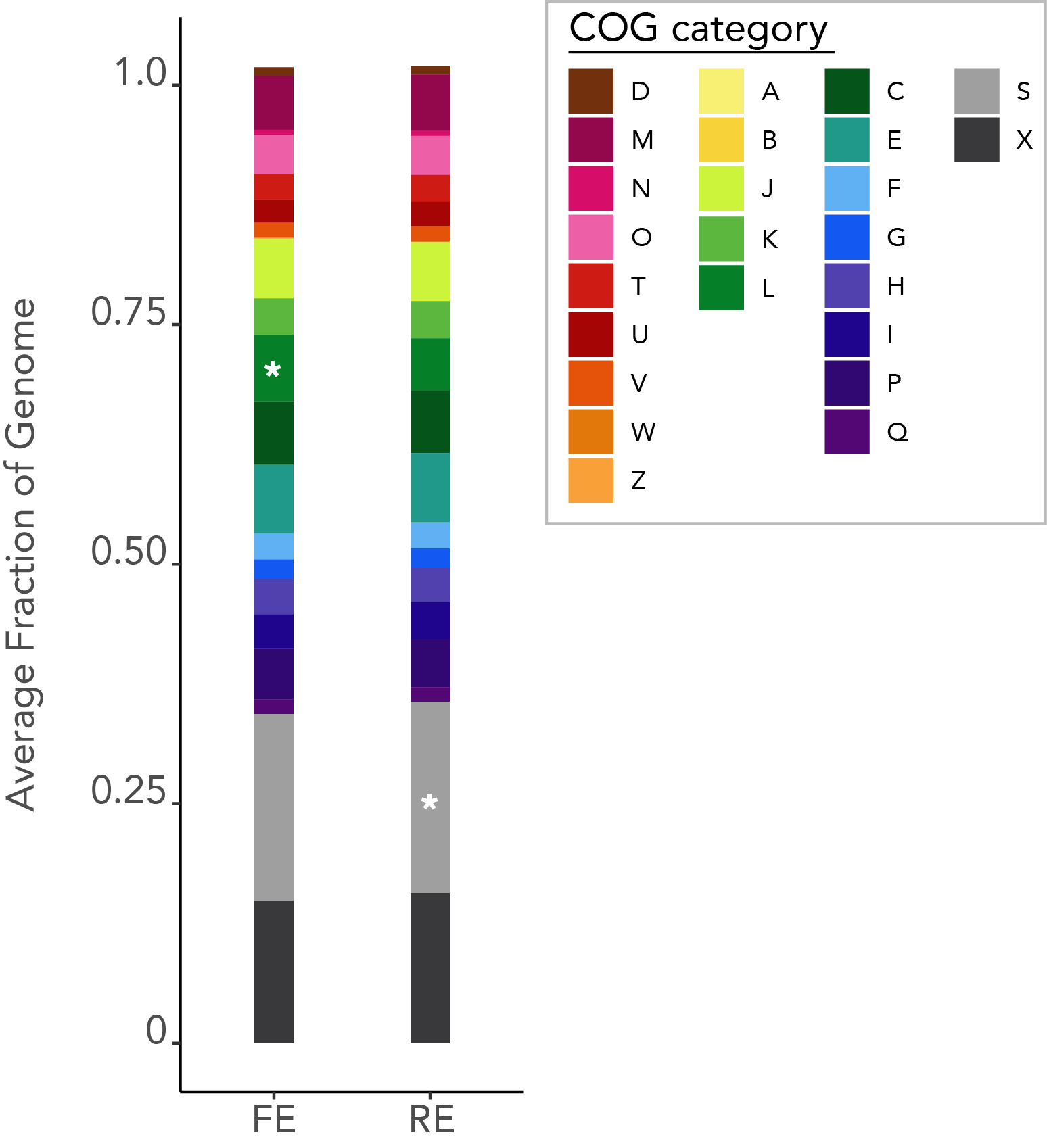

**Figure S2.** **FE and RE strains devote similar proportions of their genomes to each COG category.** Proportions of the average genome of each ecotype devoted to each cluster of orthologous genes (COG) category. Cellular processing and signaling: [D] cell cycle control cell division, chromosome partitioning; [M] cell wall/membrane/envelope biogenesis; [N] cell motility; [O] post-translational modification, protein turnover, and chaperones; [T] signal transduction mechanisms; [U] intracellular trafficking, secretion, and vesicular transport; [V] defense mechanisms; [W] extracellular structures; [Y] nuclear structure; [Z] cytoskeleton. Information storage and processing: [A] RNA processing and modification; [B] chromatin structure and dynamics; [J] translation, ribosomal structure and biogenesis; [K] transcription; [L] replication, recombination, and repair. Metabolism: [C] energy production and conversion; [E] amino acid transport and metabolism; [F] nucleotide transport and metabolism; [G] carbohydrate transport and metabolism; [H] coenzyme transport and metabolism; [I] lipid transport and metabolism; [P] inorganic ion transport and metabolism; [Q] secondary metabolites biosynthesis, transport, and catabolism. Poorly Characterized: [S] function unknown, [X] not in COG database. White asterisks indicate the COG categories that are enriched in one ecotype over the other.
